# Non-conserved metabolic regulation by LKB1 distinguishes human and mouse lung adenocarcinoma

**DOI:** 10.1101/2021.10.01.462764

**Authors:** Benjamin D. Stein, John R. Ferrarone, Eric E. Gardner, Jae Won Chang, David Wu, Qiuying Chen, Pablo E. Hollstein, Min Yuan, Roger J. Liang, John S. Coukos, Miriam Sindelar, Bryan Ngo, Steven S. Gross, Reuben J. Shaw, John M. Asara, Raymond E. Moellering, Harold Varmus, Lewis C. Cantley

**Affiliations:** Sandra and Edward Meyer Cancer Center, Weill Cornell Medicine, New York, NY 10065, USA; Department of Chemistry, University of Chicago, Chicago, IL 60637, USA; Department of Pharmacology, Weill Cornell Medicine, New York, NY 10065, USA; Molecular and Cell Biology Laboratory, The Salk Institute for Biological Studies, La Jolla, CA 92037, USA; Mass Spectrometry Core, Beth Israel Deaconess Medical Center, Boston, MA 02215, USA; Emory University, Atlanta, GA 30322, USA; Amgen, Thousand Oaks, CA 07004, USA; University of Texas – Southwestern, Dallas, TX 75390, USA; Washington University in St. Louis, Saint Louis, MO 63130, USA; Memorial Sloan Kettering Cancer Center, New York, NY 10021, USA

## Abstract

*KRAS* is the most frequently mutated oncogene in human lung adenocarcinomas (hLUAD) and activating mutations in KRAS frequently co-occur with loss-of-function mutations in the tumor suppressor genes, *TP53* or *STK11/LKB1*. However, mutation of all three genes is rarely observed in hLUAD, even though engineered mutations of all three genes produces a highly aggressive lung adenocarcinoma in mice (mLUAD). Here we provide an explanation of this difference between hLUAD and mLUAD by uncovering an evolutionary divergence in regulation of the glycolytic enzyme triosephosphate isomerase (TPI1). Using KRAS/TP53 mutant hLUAD cell lines, we show that TPI1 enzymatic activity can be altered via phosphorylation at Ser21 by the Salt Inducible Kinases (SIKs) in an LKB1-dependent manner; this allows modulation of glycolytic flux between completion of glycolysis and production of glycerol lipids. This metabolic flexibility appears to be critical in rapidly growing cells with KRAS and TP53 mutations, explaining why loss of LKB1 creates a metabolic liability in these tumors. In mice, the amino acid at position 21 of TPI1 is a Cys residue which can be oxidized to alter TPI1 activity, allowing regulation of glycolytic flux balance without a need for SIK kinases or LKB1. Our findings reveal an unexpected role for TPI1 in metabolic reprogramming and suggest that LKB1 and SIK family kinases are potential targets for treating KRAS/TP53 mutant hLUAD. Our data also provide a cautionary example of the limits of genetically engineered murine models as tools to study human diseases such as cancers.

---

Lung cancer remains the most common cause of cancer mortality in the United States and worldwide, due to high incidence coupled with poor response to standard-of-care therapies in most patients.^1^ Metabolic reprogramming is a cancer hallmark, required to support tumorigenesis in diverse environments.^2,3^ Despite improvements in our understanding of metabolic discrepancies between normal and oncogenic tissues, accurately modeling and exploiting these differences for therapeutic intervention has achieved only marginal success.

Inherent differences between humans and mice may have significant effects on tumor development through divergent mechanisms of response to the oxidative environment and to metabolic determinants.^4^ The nature and extent of such differences are unknown, but their mechanisms may illuminate unidentified molecular targets for therapy. Therefore, we sought to identify differences between human and mouse lung adenocarcinomas (hLUAD and mLUAD) with the most common genotype, mutated *KRAS* and *TP53* (KP-mutant) and determine the effects of loss of the tumor suppressor, *LKB1*, on metabolic regulation and the growth of such tumors.

## Co-occurrence of *KRAS, TP53* and *LKB1* mutations differentially affects growth of human and mouse LUADs

We used the TCGA PanCancer Atlas to determine the frequency of co-occurrence of mutations in the three most commonly mutated genes in hLUAD - *KRAS, TP53* and *LKB1* - and found that only 8 of 511 tumors carried mutations in all three genes (**Figure 1A**). A Fisher’s Exact test showed that the co-occurrence of *LKB1* and *TP53* mutations in hLUADs with a *KRAS* mutation was less frequent than expected by chance, based on the overall frequency of mutations in these three genes, with an odds ratio of 0.35 and a P-value of 0.01 (**Figure 1B**). No similar reduction was observed in the co-mutation of *TP53* and *LKB1* in the absence of *KRAS* mutations (odds ratio = 0.95; p-value of 0.87) (**Figure 1B**). A second data set from Memorial Sloan Kettering Cancer Center consisting of 1,357 lung cancer patients revealed similar exclusivity of triple mutant cases (**Figure S1A and S1B**).^5^

**Figure 1.**
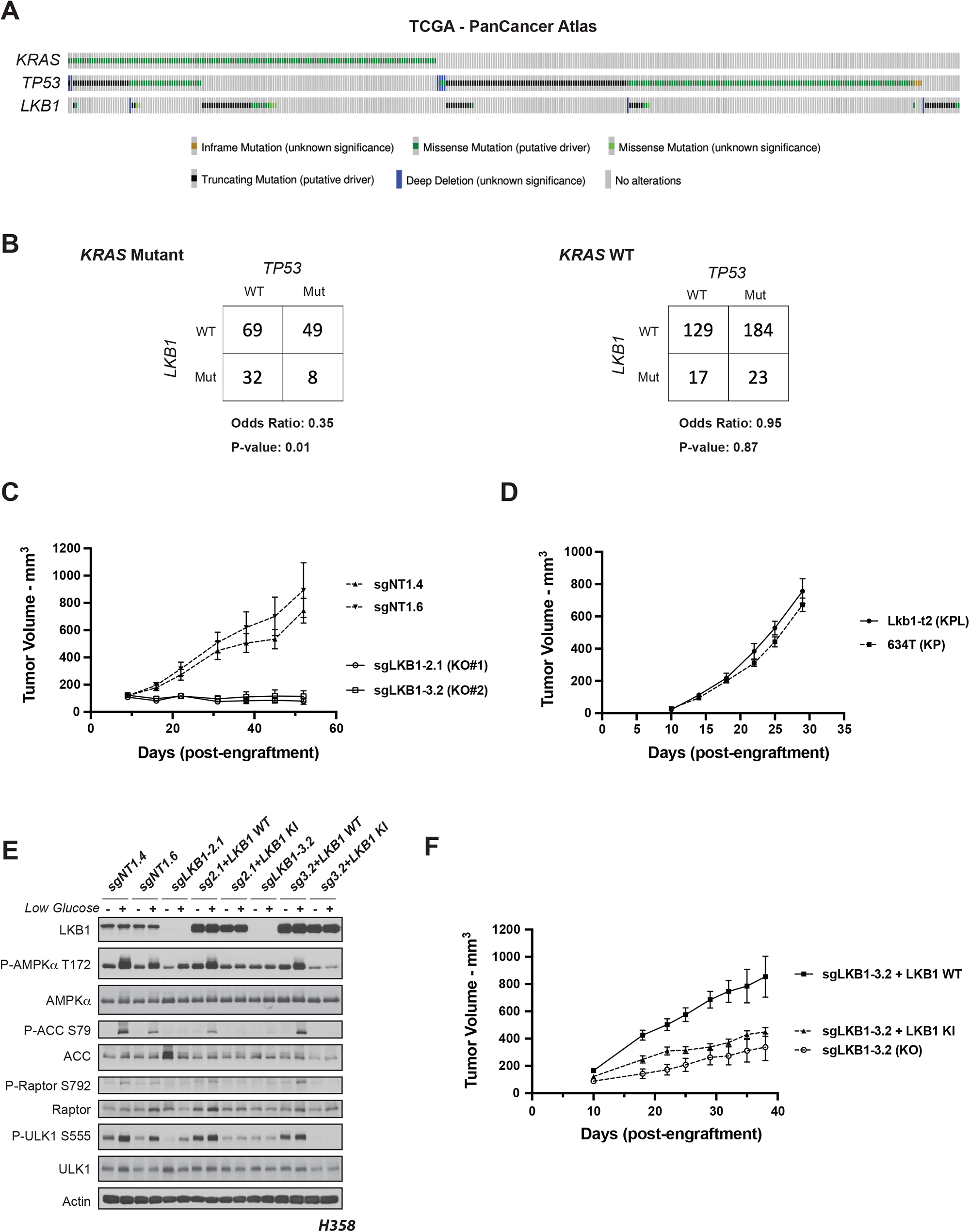
Co-occurrence of *KRAS, TP53* and *LKB1* mutations differentially affects growth of human and mouse LUADs. **(A)** The Cancer Genome Atlas PanCancer Atlas oncoprint of co-occurrence of *KRAS, TP53* and *LKB1* in human lung adenocarcinoma patients. **(B)** Fisher’s exact test of statistical likelihood of co-occurrence of *LKB1* and *TP53* mutations in a *KRAS* mutant or wildtype background respectively. **(C)** Graph of mean (+/- s.e.m.) tumor volumes of sub-cutaneous flank injections of H358 (*KRAS;TP53*) isogenic clones expressing Cas9 and a non-targeting (sgNT1.4 and sgNT1.6) or LKB1-specific (sgLKB1-2.1 and sgLKB13.2) guide RNA. 1 × 10^6^ cells implanted in right hind flank (n = 10 per cohort). **(D)** Mean (-/+ s.e.m.) volumes of mouse 634T (KP) and Lkb1-t2 (KPL) lung adenocarcinoma allograft tumors. 1 × 10^4^ cells implanted in right hind flank (*n* = 10 per cohort). **(E)** Western blot analysis of H358 (*KRAS*;*TP53*) isogenic clones (KP: sgNT1.4 and sgNT1.6; KPL: sgLKB1-2.1 and sgLKB1-3.2) and KPL lines with additional transgenic expression of guide RNA resistant LKB1 wildtype (WT) (sgLKB1-2.1 + LKB1 WT and sgLKB1-3.2 + LKB1 WT) or LKB1 kinase inactive (KI) (sgLKB1-2.1 + LKB1 KI and sgLKB1-3.2 LKB1 KI) and treated with 11.1 mM or 0.5 mM glucose for 6 hours as indicated. Restoration of AMPK signaling in LKB1 WT lines in response to 0.5 mM glucose validated by blotting for P-AMPK Thr172 and downstream substrates (P-ACC S79, P-ULK1 S555, P-Raptor S792). Similar results observed in three independent experiments and in an additional *KRAS*;*TP53* cell line, H2009 (Figure S1E). **(F)** Graph of mean (-/+ s.e.m.) tumor volumes of sub-cutaneous flank injections of H358 (*KRAS;TP53*) isogenic clones with transgenic expression of an empty vector (KO) or guide RNA resistant LKB1 wildtype (LKB1 WT) or LKB1 kinase inactive (LKB1 KI). 1 × 10^6^ cells implanted in right hind flank (*n* = 10 per cohort).

While mutations in *KRAS, TP53* and *LKB1* together are rare in hLUAD, previous studies have shown that genetically engineered mouse models (GEMMs) harboring conditional mutations in all three genes develop mLUAD that is more aggressive and more likely to metastasize than mLUAD with only two of these genes mutated.^6–9^ To investigate this discrepancy between LUAD in human patients and mouse models, we first generated isogenic clones of human KP cell lines with and without LKB1 deficiency and compared them to existing GEMM-derived mouse tumor lines with parallel genotypes.^8^ The human KP lines engrafted and formed tumors *in vivo*, whereas isogenic lines in which *LKB1* was deleted (KPL) did not (**Figure 1C**). Furthermore, human KP lines readily formed spheroids in organotypic culture, but KPL lines did not (**Figure S1C**). In contrast, GEMM-derived KP and KPL lines both formed tumors *in vivo* and spheroids *in vitro* (**Figures 1D and S1D**).

To determine whether these observations were attributable to LKB1 kinase activity, wildtype or kinase-inactive (K78I) LKB1 were re-expressed in isogenic KPL hLUAD lines derived from two human KP lines. We first verified that wildtype LKB1 restored the activity of AMP-activated protein kinase (AMPK), a known substrate of LKB1, under conditions of energy stress. Glucose restriction caused LKB1-dependent phosphorylation of AMPK at Thr172 and of its downstream substrates (Acetyl CoA Carboxylase (ACC) at Ser79, Raptor at Ser792, and Unc-51 Like autophagy activating Kinase 1 (ULK1) at Ser555) (**Figure 1E and Figure S1E**). Expression of wildtype (WT), but not kinase-inactive (KI), LKB1 rescued growth of the xenografts in immunodeficient mice, suggesting that LKB1 kinase activity is required to support tumor formation by human KP LUAD cells (**Figure 1F**).

### Phosphorylation of human TPI1 is LKB1-dependent

Since LKB1 phosphorylates and activates a family of AMPK-related Ser/Thr protein kinases (AMPKRs) involved in regulating various metabolic and stress response pathways, we used comprehensive quantitative phospho-proteomics under glucose-limited conditions to assess differences in protein phosphorylation between KP and KPL isogenic human lines. Phosphorylation of Ser21 on the glycolytic enzyme Triosephosphate Isomerase (TPI1) was one of the most significantly down-regulated phosphorylation events observed when comparing KPL to KP (**Figure 2A, S2A** and **S2B**). As expected, we also observed reduced phosphorylation of Ser108 in the beta subunits of AMPK: PRKAB1, and PRKAB2, each required for enzymatic activity (**Figures S2B**). In contrast, using the same experimental design with tumor-derived mouse cell lines, we did not detect phosphorylation of Tpi1 in cells with either KP or KPL genotypes (**Figure S2C**).

**Figure 2.**
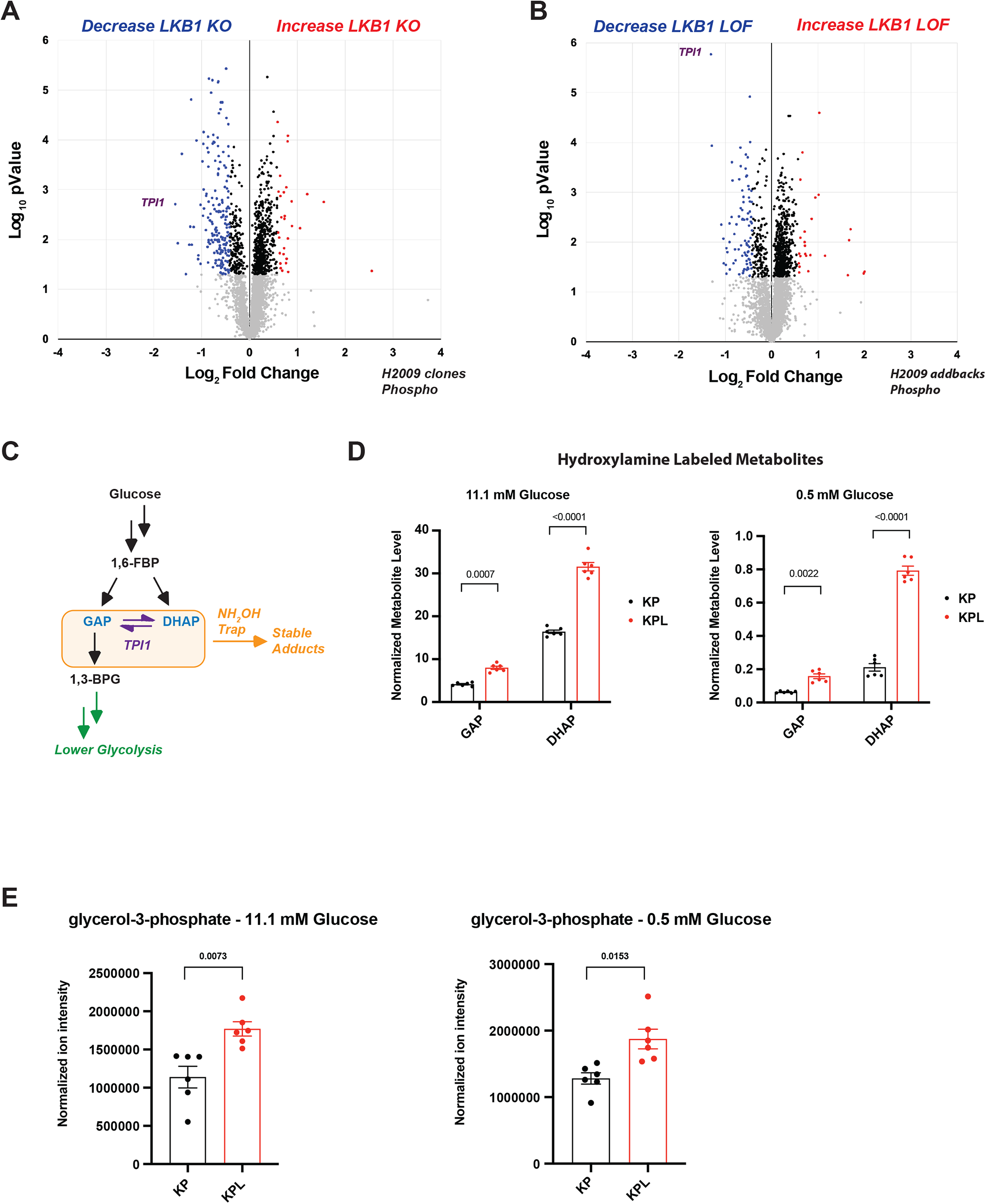
Phosphorylation of human TPI1 is LKB1-dependent and regulates triose phosphate levels. **(A)** Volcano plot of quantitative phospho-proteomic data of genetic sensitivity in H2009 clones (2 KP clones and 2 KPL clones), two biological replicates each, N = 4 per genotype. Cells grown in 0.5 mM glucose for 6 hours. Phospho-peptides that pass statistical criteria (p-value < 0.05) are highlighted in black, red and blue, peptides that do not satisfy this are colored grey. Phospho-peptides colored red satisfy a fold change > 1.5; colored blue, fold change < −1.5. TPI1 P-Ser21 peptide labeled in purple text. **(B)** Volcano plot of quantitative phospho-proteomic data of genetic sensitivity in H2009 isogenic clones including clones with transgenic expression of guide RNA resistant wildtype (WT) or kinase inactive (KI) LKB1 in LKB1-specific knockouts (sgLKB1-3.1 and sgLKB1-3.7) from Figure S1E, 4 biological replicates each. LKB1 Loss-of-function (LOF) group consisted of merging LKB1 knockout lines (KPL: sgLKB1-3.1 and sgLKB1-3.7) with lines expressing guide RNA resistant LKB1 KI (KPL + LKB1 KI: sgLKB1-3.1 + LKB1 KI and sgLKB1-3.7 + LKB1 KI); and compared to H2009 lines containing non-targeting guide RNA (KP: sgNT1.1 and sgNT1.2) merged with LKB1 knockout lines expressing guide RNA resistant LKB1 WT (KPL + LKB1 WT: sgLKB1-3.1 + LKB1 WT and sgLKB1-3.7 + LKB1 WT) at the phospho-peptide level. Cells were grown in 0.5mM glucose for 6 hours. Statistical criteria and color scheme same as for panel A. TPI1 P-Ser21 peptide labeled in purple text. **(C)** Schematic showing metabolites (shaded in the orange box) chemically labeled to create stable adducts. **(D)** *In-situ* chemical trapping metabolomics of hydroxylamine-labeled GAP and DHAP in H2009 clones (KP: sgNT1.1 and sgNT1.2; KPL: sgLKB1-3.1 and sgLKB1-3.7) treated in culture for 6 hours with 11.1 mM or 0.5 mM respectively. Data presented are representative of three independent biological experiments each containing three technical replicates and reported as the mean (-/+s.e.m.). Cell number normalized across models 12 hours prior to assay and samples normalized to an exogenous standard, *d* _3_-serine. Statistical significance determined by two-tailed paired t-test. **(E)** Normalized ion intensity of glycerol-3-phosphate from steady-state analysis of H2009 clones treated for 30 minutes with 11.1 or 0.5 mM glucose. Analysis conducted in H2009 isogenic clones (KP: sgNT1.1 and sgNT1.2; KPL: sgLKB1-3.1 and sgLKB1-3.7) in biological triplicate and reported as the mean (-/+s.e.m.). Statistical significance determined by two-tailed paired t-test.

To assess if restoration of LKB1 kinase activity re-established phosphorylation of TPI1 during metabolic stress, we used quantitative proteomics and phospho-proteomics to analyze human KPL cells expressing WT or KI LKB1 in parallel with KP and KPL cells under glucose-limited conditions. Ser21 phosphorylation (p-Ser21) on TPI1 was again one of the most significantly reduced phosphorylation sites when KPL cells and KPL cells expressing KI LKB1 were compared with KP cells and KPL cells expressing WT LKB1 (**Figure 2B**). Furthermore, quantification of phosphopeptide ion intensities within individual genotypes confirmed restoration of p-Ser21 levels in human KPL lines expressing WT, but not KI LKB1 (**Figure S2D**), without significant variation in the abundance of TPI1 protein.

### Phosphorylation of TPI1 regulates triose phosphate levels

To examine the possibility that loss of regulation of TPI1 in hLUAD might explain selection against the KPL genotype, we studied the metabolic consequences of LKB1 deficiency. It is known that TPI1 controls the interconversion of the triose phosphates, dihydroxyacetone phosphate (DHAP) and glyceraldehyde-3-phosphate (GAP), both of which are generated from the upstream glycolytic intermediate fructose-1,6-bisphosphate (1,6-FBP) by aldolase. This conversion in carbon metabolism lies at a critical bifurcation point: one product, GAP, is used for glycolysis and energy homeostasis, whereas the other, DHAP, is used for lipid synthesis, cellular growth, and has recently been shown to activate the mammalian Target of Rapamycin protein kinase (mTOR).^10^ Additionally, previous studies have shown that increased oxidative burden due to *KRAS* and/or *TP53* mutations cause metabolic flux to primarily flow through the oxidative Pentose Phosphate Pathway (oxPPP) to increase reductive potential and restore redox balance to overcome this liability.^11–13^

We next assessed the influence of LKB1-dependent phosphorylation on TPI1 activity by measuring the pools of GAP and DHAP in KP and KPL hLUAD cell lines. Due to the inherent instability and complex chromatographic separation of the triose phosphates, we used *in situ* chemical-trapping metabolomics with hydroxylamine labeling of live cells under normal and glucose limited conditions prior to lysis to create stable adducts and measured them (**Figure 2C and S2E**).^14^ These analyses confirmed relative elevation of DHAP in KPL lines, further suggesting that TPI1 phosphorylation limited DHAP accumulation to maintain GAP for glycolysis, crucial under glucose-limited conditions (**Figure 2D**). Additionally, a parallel analysis including KPL cell lines expressing WT and KI LKB1 with the same method revealed that levels of GAP and DHAP in WT LKB1 wildtype lines more accurately recapitulated endogenous levels in KP human cells, while DHAP remained elevated in the LKB1-KI cells (**Figure S2F**). Furthermore, steady-state analysis revealed that human KPL cells had a significant increase in glycerol-3-phosphate, the next metabolic intermediate in the lipid and triglyceride synthesis pathway under normal and low glucose conditions (**Figure 2E**). Collectively, the observed changes in metabolites and phosphorylation of human TPI1 suggested that LKB1 regulates distribution of glycolytic metabolites via the triose phosphates through a regulatory phosphorylation site in TPI1.

### Non-conserved amino acid sequence of TPI1 requires LKB1 to regulate its multimeric state in human but not mouse LUAD

To determine whether differences between LKB1 loss in human and mouse LUAD cells could be explained by differences in regulation of TPI1/Tpi1, we explored the evolutionary conservation of the primary amino acid sequence surrounding position 21 of this enzyme. Ser21 and the surrounding residues are conserved in most mammals and many other metazoan organisms, including yeast. However, in mouse and rat Tpi1, Ser21 has been replaced by a cysteine (**Figure 3A**). In a published crystal structure of human TPI1, the hydroxyl moiety of Ser21 is located at a region of subunit:subunit interactions, stabilizing the homodimer. Notably, the nearby residues Arg18 and Lys19 form inter-subunit electrostatic interactions predicted to further stabilize the highly active homodimeric state. This structure raises the possibility that phosphorylation of Ser21 could produce intra-subunit charge interactions with Arg18 and/or Lys19, interfering with the ability of these amino acids to confer stability to the dimer, thereby altering enzymatic activity (**Figure 3B**).^15^ Since the sulfur atom of Cys21 in rodent Tpi1 could be oxidized to sulfinic or sulfonic acid, mimicking phosphorylation of Ser21, it is possible that rodents have a mechanistic alternative to phospho-dependent regulation of TPI1 activity. This could explain differences in the response to loss of LKB1 in mouse and human tumors, circumventing the requirement for LKB1 activity in murine tumors.

**Figure 3.**
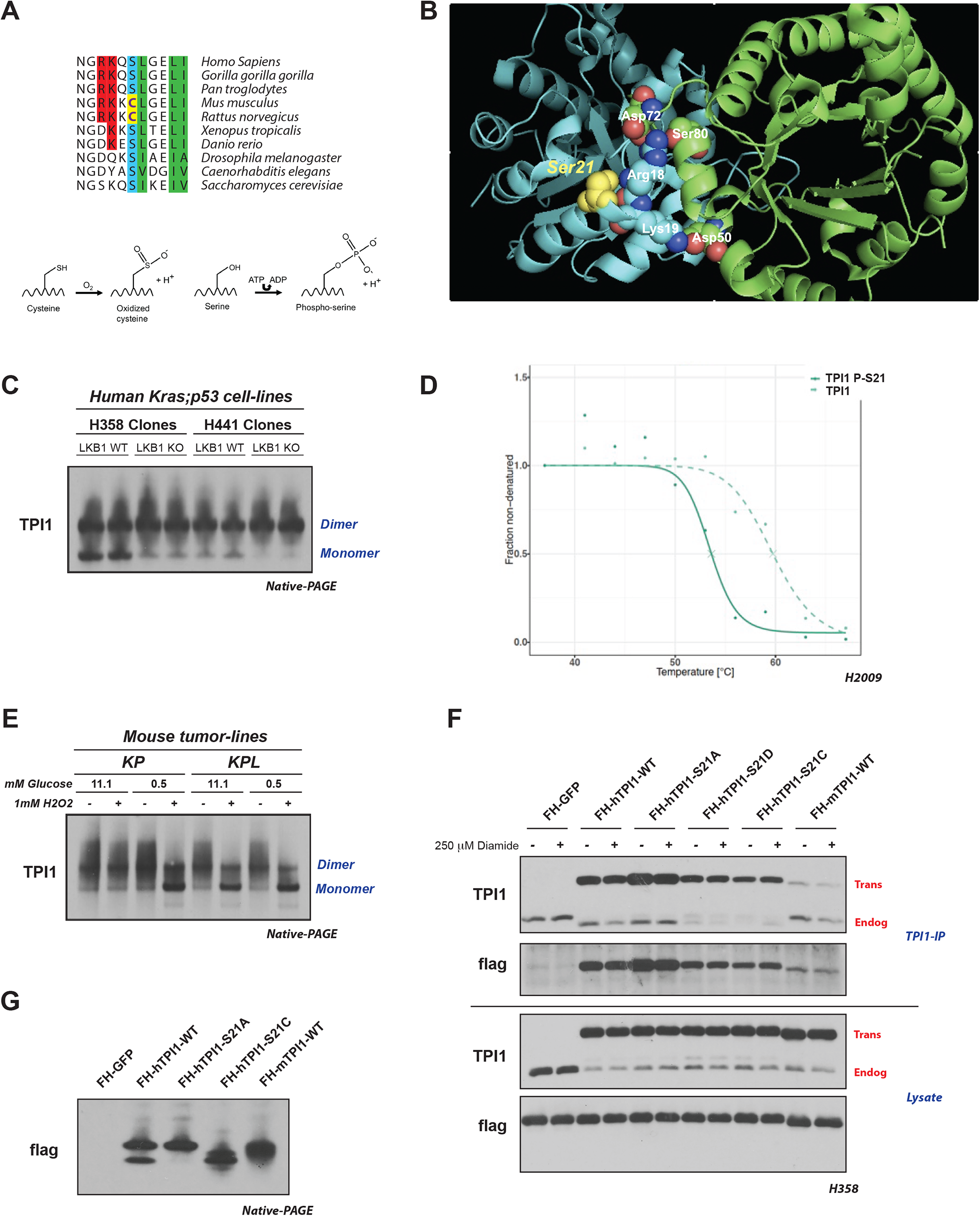
LKB1 regulates the multimeric state of TPI1 in KP-mutant hLUAD but not in mLUAD cell lines due to an amino acid difference at position 21. **(A)** Sequence alignment of TPI1 amino acid residues 16 to 26 across species, showing conservation of Ser21 from H. sapiens to *S. Cerevisiae*, with cysteine at position 21 in mouse and rat Tpi1. Cartoon comparing predicted side-chain chemistry, with oxidized cysteine and phosphorylated serine, is drawn below. **(B)** Crystal structure of TPI1 homodimer (cyan and green respectively) with critical residues highlighted in space-filling atoms. Serine 21 on the cyan monomer is highlighted in yellow. **(C)** Western blot analysis of Blue Native PAGE of human isogenic clones derived from H358 and H441 cell lines. Cells were grown under normal conditions (11.1 mM glucose). **(D)** Melting curve plot from Thermal Proteome Profiling of unmodified and Serine 21 phosphorylated TPI1. Analysis conducted in H2009 isogenic clones expressing Cas9 and a non-targeting (sgNT1.1 and sgNT1.2) or LKB1-specific (sgLKB1-3.1 and sgLKB13.7) guide RNA. **(E)** Western blot (Blue Native PAGE) of extracts from mLUAD cell lines. Cells were cultured in either 11.1 mM or 0.5 mM glucose for 6 hours then treated with 1 mM H_2_O_2_ for 15 minutes. **(F)** Western blot of proteins co-immunoprecipitated from extracts of H358 cells expressing Cas9 and a non-targeting (FH-GFP cell line) or TPI1-specific (all other cell lines) guide RNA and transgenic expression of Flag-HA tagged GFP or guide RNA resistant TPI1 allelic variants using a polyclonal antibody against full-length TPI1. Cells were cultured in 11.1 mM glucose and treated with 250 μM Diamide or vehicle for 15 minutes prior to collection. **(G)** Western blot (Blue Native PAGE) of extracts of H358 cell lines used for co-immunoprecipitation in panel F. Cells were cultured in 11.1 mM glucose.

Based on the structural features of human and mouse TPI1/Tpi1, we next asked whether the loss of LKB1 kinase activity, which prevents phosphorylation of human TPI1 at Ser21, would differentially affect the dimerization and activity of this enzyme in cells from the two species. We used native gel electrophoresis (BN-PAGE) and western blotting to determine the proportions of monomeric and dimeric TPI1 in extracts of two human KP cell lines, in the presence and absence of LKB1, when cells are grown under normal glucose conditions (**Figure 3C**). Loss of LKB1 promoted the dimeric (more slowly migrating) form of the human protein; conversely, cell lines expressing LKB1 had increased monomeric (more rapidly migrating) TPI1 (Figure 3C). Thermal Proteome Profiling (TPP) was also used to measure the thermal stability of TPI1 proteoforms.^16,17^ The ΔTm (measured at 0.5 fraction denatured) of the phosphorylated variant was 5.8°C lower than that of unmodified TPI1, further supporting the prediction that phosphorylation of Ser21 disrupts TPI1 dimerization (**Figure 3D**).

In contrast, we observed no changes in the ratio of the monomeric and dimeric forms of mouse Tpi1 in KP versus KPL mLUAD cell lines at high (11.1 mM) or low (0.5 mM) glucose concentrations (**Figure 3E)**. However, acute treatment of the mouse lines with the oxidant peroxide caused a dramatic shift towards the rapidly migrating (monomeric) form of Tpi1 in low glucose medium, regardless of Lkb1 status (Figure 3E). Additionally, peroxide treatment caused a similar shift towards monomeric Tpi1 under high (11.1 mM) glucose conditions in KPL, but not KP mouse cells (Figure 3E), consistent with earlier reports that loss of Lkb1 increases basal oxidative stress.^18–21^

To further explore the functional significance of the amino acid difference at position 21 of TPI1 and Tpi1 and its effect on homodimer formation, we created an allelic panel of FLAG-HA-tagged TPI1 variants expressed as transgenes in human KP cells following deletion of endogenous TPI1. We observed that replacement of Serine with Alanine (S21A) increased recovery of both the transgenic and remaining endogenous TPI1 by immunoprecipitation, implying that an inability to phosphorylate position 21 of TPI1 stabilized the TPI1 dimer. In contrast, the phospho-mimetic S21D mutant form of TPI1 or a mutant in which Ser21 is replaced by an oxidizable cysteine (S21C) significantly reduced co-immunoprecipitation of the remaining endogenous TPI1 (**Figure 3F**). These findings were further confirmed when TPI1 variants from the allelic panel were analyzed by BN-PAGE. Wildtype transgenic human TPI1 was found in both the dimeric and monomeric states, but the S21A mutant was detected solely in the dimeric state, and the S21C variant was mostly monomeric (**Figure 3G**). Collectively, these findings support the conclusion that phosphorylation of human TPI1 or oxidation of murine Tpi1 destabilizes its dimeric form, providing a structural mechanism by which TPI1/Tpi1 activity can be regulated in response to LKB1-dependent phosphorylation or by oxidative stress.

### LKB1-activated members of the Salt Inducible Kinase family phosphorylate human TPI1

We next sought to determine whether human TPI1 is phosphorylated directly by LKB1 or by one of the downstream LKB1-dependent Ser/Thr protein kinases of the AMPKR family; these kinases are known to mediate responses to various metabolic stresses, all of which require phosphorylation by LKB1 for activity (**Figure 4A**).^22–24^

**Figure 4.**
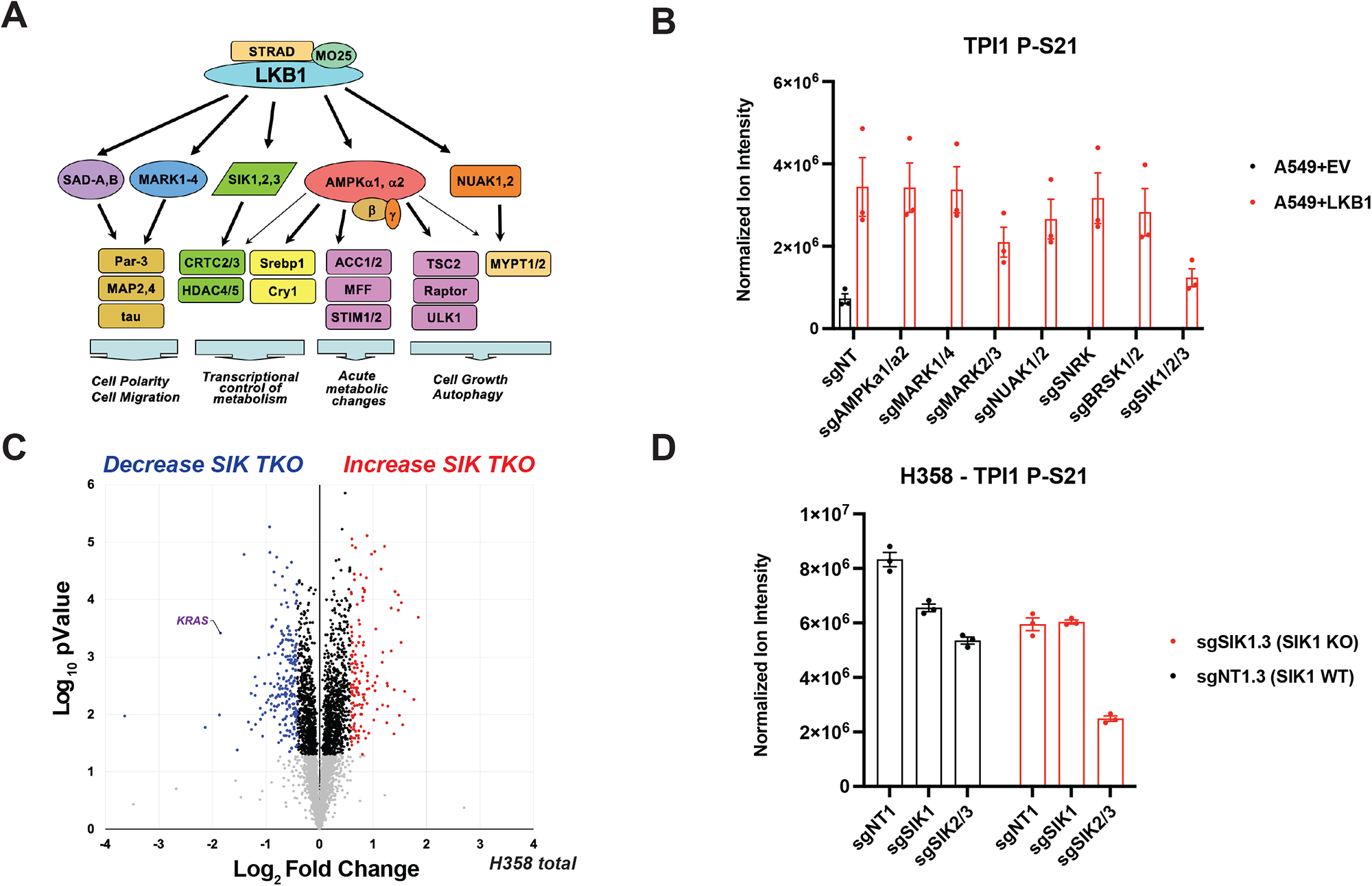
Salt Inducible Kinases phosphorylate human TPI1 in KP hLUAD cell lines. **(A)** Cartoon depicting regulation of the AMPK-related (AMPKR) kinase family members by LKB1 and their downstream substrates. **(B)** Bar graph of normalized ion abundance for the TPI1-derived ser-21 phospho-peptide from extracts of A549 cell-lines infected with an empty vector or a vector expressing wild type LKB1; the indicated guide RNAs were used to inactivate members of the AMPKR subfamilies. Cell lines were cultured in 11.1 mM glucose prior to analysis. Ion intensities were normalized to identified non-phosphorylated variants across conditions to control for protein expression and reported as the mean (-/+ s.e.m.). **(C)** Volcano plot of quantitative proteomic data used to compare protein expression in clones of H358 (2 KP clones and 2 KP SIK TKO clones, with 2 biological replicates of each). Cells were cultured in 11.1 mM glucose for 6 hours before lysis. Proteins that pass statistical criteria (p-value <0.05) are highlighted in black, red and blue; those that do not satisfy this criterion are colored grey. Proteins highlighted in red satisfy the fold change threshold (>1.5) after triple deletion of SIK1,2,3. Proteins highlighted in blue satisfy the fold change threshold of < −1.5) for a decrease after SIK1,2,3 triple deletion. KRAS is labeled in purple text. **(D)** Bar graph of normalized ion abundance for the TPI1-derived, ser-21 phospho-peptide in extracts of isogenic H358 cell-lines containing a non-targeting control (sgNT1.3) or SIK1 specific (sgSIK1.3) guide RNA and additional control (NT1) SIK1 (sgSIK1) or dual SIK2 and SIK3 (sgSIK2/3) guide RNAs. Ion intensities were normalized against identified non-phosphorylated variant across conditions. Cell lines were cultured in 11.1 mM glucose prior to lysis, analyzed in biological triplicate, and reported as the mean (-/+ s.e.m.).

To determine whether AMPKR kinases are directly responsible for TPI1 phosphorylation downstream of LKB1, we monitored phosphorylation of Ser21 in TPI1 in a panel of human *KRAS;LKB1*-mutant cell lines in which sub-families of the AMPKR kinases have been genetically eliminated, after restoring stable expression of WT LKB1 from a transgene (**Figure 4B**).^7^ Restoration of LKB1 increased the phosphorylation of Ser21 in TPI1, consistent with results in Figure 2B and S2D, and deletion of the Salt Inducible Kinase (SIK) subfamily significantly reduced Ser21 phosphorylation (**Figure 4B**). Deletion of other AMPKR super-family members [the Microtubule Affinity Regulating Kinases (MARKs), the NUAK Family Kinases (NUAKs), the Brain-Specific Serine/Threonine-Protein Kinases (BRSKs), the catalytic subunits of AMPK, and SNF Related Kinase (SNRK)] did not have significant effects on Ser21 phosphorylation of TPI1. Furthermore, by analyzing specific combinations of deletions of SIK family members, we found that SIK1 and SIK3 together made the greatest contribution to phosphorylation of Ser21 in TPI1 **(Figure S3A**).

In agreement with the concept that the SIK sub-family of protein kinases drive phosphorylation of Ser21 in TPI1, phosphorylation of Ser551 in SIK3, known to regulate activity through altering molecular association, was one of the most significantly down-regulated phosphorylation sites in the LKB1-deleted, KP-mutant hLUAD cell lines (Figure S2A).^25,26^ Additionally, phosphorylation of the regulatory sites in a SIK-family substrate, CREB Regulated Transcription Coactivator 3 (CRTC3), was also down-regulated in KPL versus KP human lines (Figures S2A and S2B). Furthermore, we found that the amount of *SIK1* mRNA was significantly increased upon inactivation of *LKB1* in multiple human KP lines, suggesting that a signaling network might increase transcription of *SIK1* via a feedback mechanism to recover SIK activity after loss of LKB1 (**Figure S3B**).

We next asked if SIK family kinases were responsible for phosphorylation of TPI1 in human KP lines that express LKB1 from the endogenous *LKB1* locus. We generated a series of cell lines deficient in members of the *SIK* gene family, including two *SIK1/2/3* triple knockout lines (SIK TKO). Analysis of this series of cell lines by western blot confirmed significant deletion of all SIK family members in the SIK TKO lines (**Figure S3C**). A quantitative proteomic analysis of differences between SIK WT and SIK TKO cells revealed that KRAS was among the most significantly down-regulated proteins (**Figure 4C**). This observation is consistent with SIK family enzymes being critical for cell growth when mutant *KRAS* is present and *TP53* is mutated. Measurement of the TPI1 p-Ser21 tryptic peptide from cells with various combinations of SIK1, SIK2 and SIK3 deletions argues that each of these kinases contribute to phosphorylation of TPI1 at Ser21 in human KP cells **(Figure 4D**).

Taken together, these results indicate that in the KP hLUAD cell lines investigated, the SIK family of LKB1-regulated protein kinases appear to dominate the phosphorylation of Ser21 in TPI1, although deletion of all three SIK family members did not eliminate phosphorylation at this site, suggesting that other LKB1-regulated protein kinase may also contribute to TPI1 phosphorylation or compensate for loss of the SIK kinases. Our results suggest that an LKB1 inhibitor might be an effective therapy for KP mutant hLUADs, though there are likely to be significant toxicities associated with the inhibition of LKB1. An alternative, potentially less toxic therapy would inhibit a sub-group of LKB1-regulated protein kinases, including the SIK family kinases, in *KRAS*;*TP53* human lung adenocarcinoma.

## Discussion

Although much progress has been made using GEMMs to decipher mechanisms of tumor initiation and progression, comparisons of human and mouse tumors often lead to conflicting observations.^27^ In particular, accurately recapitulating the tumor metabolic environment remains a significant challenge, but an important one, since discrepancies between mouse and human tumors are likely to have implications in development of novel therapeutic agents.^28^ Here we provide a mechanistic explanation for why the loss of LKB1 in hLUADs driven by *KRAS* and *TP53* mutations is a rare event that appears to be selected against in human tumors, but not in mouse tumors, where the loss of Lkb1 enhances tumorigenesis and metastasis.

Several recent reports have implicated the SIK kinases as effectors of LKB1-mediated tumor suppression in *Kras*- and *Kras*;*Tp53*-mutant mLUAD; however similar findings have not been reported in hLUADs.^7,29^ Here we find that this discrepancy may be due, at least in part, to a single amino acid difference between rodent and other metazoan versions of the glycolytic enzyme TPI1. In turn, that difference can influence subsequent metabolic events, that determine the flow of glucose-derived tri-carbon substrates into pathways for glycolysis or lipid synthesis. In humans, the abundance of the products of TPI1 is governed by the LKB1-SIK-TPI1 signaling axis that we have elucidated in this manuscript. In rodents, the substitution of an oxidizable cysteine for a phosphorylatable serine at residue 21 of Tpi1 enables direct redox regulation, circumventing the requirement for regulation by LKB1-SIK-mediated phosphorylation. Our biochemical, proteomic, and metabolomic data support the conclusion that phosphorylation of TPI1 in hLUAD regulates the biophysical distribution of monomeric and dimeric forms, altering enzymatic activity and in turn triose phosphate pools. This reduces the conversion of GAP to DHAP, an energetically downhill reaction, and thereby shifts the balance away from glycerol lipid production and towards alternate metabolic pathways, including glycolysis and the TCA cycle. Regulation of metabolites at this central point in the glycolytic pathway could help to overcome metabolic stresses experienced during tumorigenesis and to improve the efficiency of energy production. In addition, this regulation allows rapidly growing cells to balance pathways for lipid synthesis versus serine/glycine synthesis. Collectively, these metabolic differences could strongly influence a wide range of pro-tumorigenic processes and have significant effects on tumor cell phenotype, all of which warrant future study. Additional features of these phenomena - such as how specific *KRAS* and *TP53* mutations influence this phenotype and their contributions to the response to metabolic and oxidative stresses - have yet to be deciphered.

Knowledge of the differences in human and mouse TPI1/Tpi1 may not only explain the different consequences of loss of LKB1 in human and mouse LUADs but may also help to design next-generation mouse models in which the mechanisms of metabolic regulation of human cancers are more accurately replicated. Furthermore, the research reported here suggests that selective inhibitors of LKB1 or of SIK family protein kinases could be effective in treating human Kras/TP53 mutant lung cancers or other cancers with KRAS/TP53 mutations. But our work also raises the cautionary note that preclinical trials with such inhibitors would likely fail in currently available GEMMs with Kras and Tp53 mutations.

Finally, the observations reported here also reveal new ways for LKB1 to regulate metabolism, beyond its known capacity to respond to cellular energy levels through activation of AMPK.^30^ While enzymes such as hexokinase, pyruvate kinase and phosphofructokinase have been intensely studied in regard to phosphorylation-dependent regulation in cancers, TPI1 has not been considered a likely site for cancer-dependent regulation of metabolic flux. Additional research is needed to understand how critical this regulation is to other types of human cancers and whether this knowledge can lead to new cancer therapies across multiple organ types and multiple mutational backgrounds.

## Data Availability

All derived MS/MS data will be deposited on MASSive and ProteomeXchange.

## Supplemental Information

Supplemental Information includes 3 supplemental figures.

## Acknowledgements

We thank Kwok-Kin Wong for kindly providing GEMM derived LUAD cell-lines; 634T and Lkb1-t2. This work was supported, in part, by NIH grants; P01CA120964 (to L.C.C., R.J.S. and J.M.A.), R35CA197588 (to L.C.C.), R35-CA220538 (to R.J.S.), R01-DK080425 (to R.J.S.), R01AR076029 (to Q.C.) and R21ES032347 (to Q.C.); NSF-CAREER CHE-1945442 (to R.E.M.) and the Alfred P. Sloan Foundation FG-2020-12839 (to R.E.M.). H.E.V. is the Lewis Thomas University Professor at Cornell University. J.R.F. is the Lee Cooperman Physician-Scientist of the Damon Runyon Cancer Research Foundation (DRG 18-18). E.E.G. is the Kenneth G. and Elaine A. Langone Fellow of the Damon Runyon Cancer Research Foundation (DRG-2343-18). B.N. is supported by a National Cancer Institute (NCI) of the National Institutes of Health (NIH) F99/K00 Career Transition Fellowship (F99CA234950).

## Author Contributions

B.D.S., H.E.V., and L.C.C. conceived of and designed the study. B.D.S. guided and performed most experiments, performed all proteomics and biochemical experiments, metabolomics experiments and all computational analyses. J.R.F. performed experiments and analyzed clinical data. E.E.G., D.W. and B.N. assisted in xenograft studies in Figures 1C, 1D and 1F. E.E.G. performed 3D Matrigel experiments in Figure S1C. J.W.C., J.S.C. and R.E.M. performed and analyzed chemical trapping metabolomics data in Figure 2D. M.Y. and J.M.A. performed metabolomics analyses in Figures 2E. Q.C., M.S. and S.S.G. analyzed metabolomics experiments. P.E.H. and R.J.S. provided cell-lines and lysates utilized for proteomics in Figure 4B and S4A. R.J.L. performed experiments. B.D.S. wrote the manuscript, which was reviewed by all authors. H.E.V. and L.C.C. supervised the study.

## Competing Interests

L.C.C. is a co-founder and member of the SAB and holds equity in Faeth Therapeutics, Volastra Therapeutics and Larkspur Therapeutics. He is also a co-founder, former member of the SAB and BOD and holds equity in Agios Pharmaceuticals. H.E.V. is a member of the SABs of Volastra, Dragonfly Therapeutics, and Surrozen. These companies are developing novel therapies for cancer. L.C.C.’s laboratory has previously received some financial support from Petra Pharmaceuticals. None of these companies are developing drugs related to the research in this paper. All other authors declare no competing interests.

## Supplemental Figure Legends

**Figure S1.**
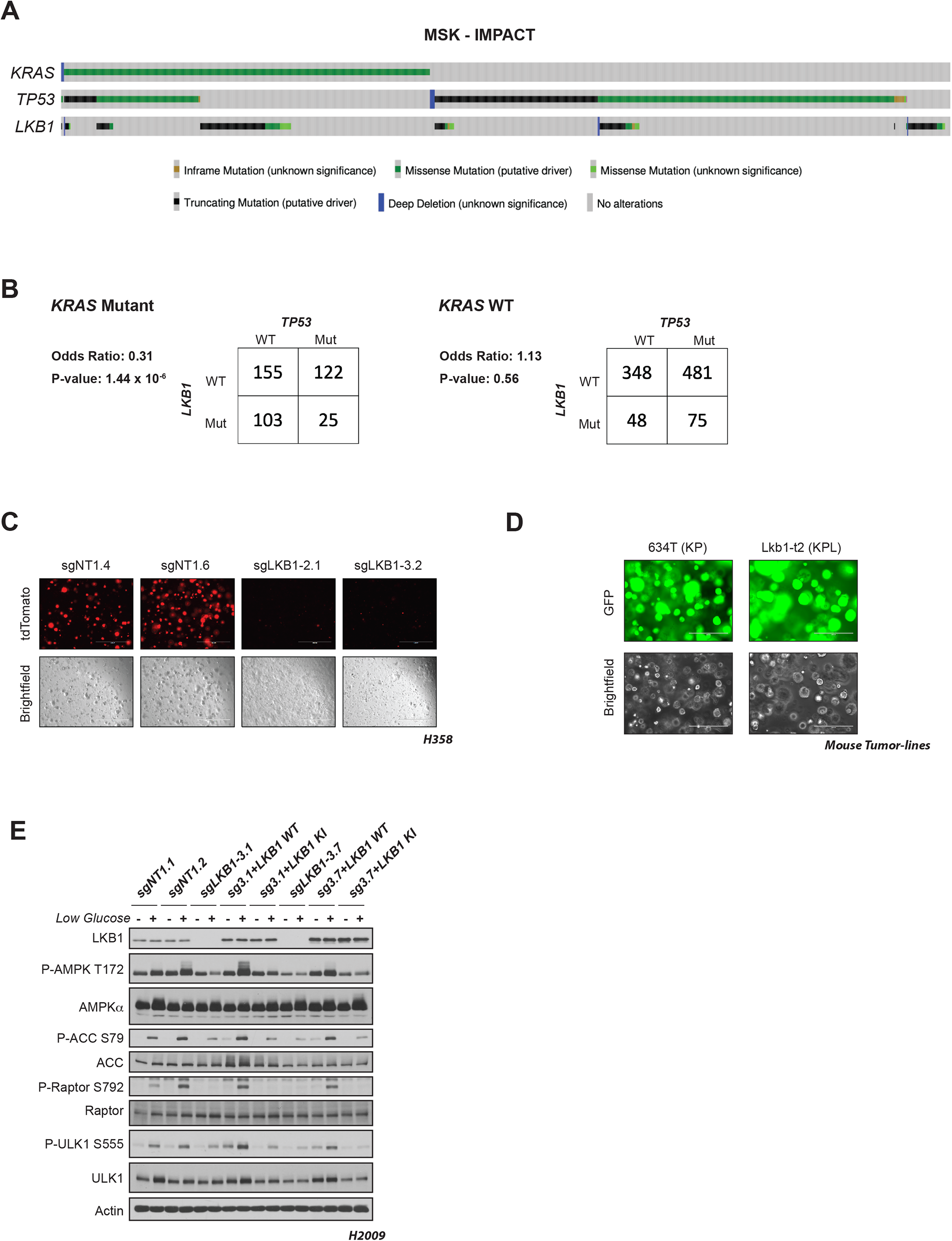
Differential co-occurrence and effects of *KRAS, TP53* and *LKB1* mutations on human and mouse LUADs. **(A)** MSK Impact oncoprint of co-occurrence of *KRAS, TP53* and *LKB1* mutations in human lung adenocarcinomas, with Fisher’s exact test of statistical likelihood of co-occurrence of *LKB1* and *TP53* mutations in a LUAD with a *KRAS* mutant or wildtype background. **(C)** 3D spheroid growth in Matrigel of isogenic clones of the H358 cell line labeled with a tdTomato fluorescent reporter and expressing CAS9 and non-targeting controls (sgNT1.4 and sgNT1.6) or LKB1-specific (sgLKB1-2.1 and sgLKB1-3.2) guide RNAs. 5,000 cells were seeded into Matrigel and grown for 10 days in media changed every 24 hours. Images taken on EVOS fluorescence microscope under 4x magnification and filter to resolve tdTomato signal intensity and brightfield. **(D)** 3D spheroid growth in Matrigel of GEMM-derived mLUAD cell lines containing transgenic lentiviral expression of GFP under control of a CMV promoter. 5,000 cells were seeded into Matrigel and assay was conducted for 10 days in culture media changed every 24 hours. Images taken on EVOS fluorescence microscope under 10x magnification and filter to resolve GFP signal intensity and brightfield. **(E)** Western blot analysis of H2009 (*KRAS*;*TP53*) isogenic clones (KP: sgNT1.1 and sgNT1.2; KPL: sgLKB1-3.1 and sgLKB1-3.7) and lines with additional transgenic expression of guide RNA resistant LKB1 wildtype (WT) (sgLKB1-3.1 + LKB1 WT and sgLKB1-3.7 + LKB1 WT) or LKB1 kinase inactive (KI) (sgLKB1-3.1 + LKB1 KI and sgLKB1-3.7 LKB1 KI) and treated with 11.1 mM or 0.5 mM glucose for 6 hours as indicated. Restoration of AMPK signaling in LKB1 WT lines in response to 0.5 mM glucose validated by blotting for P-AMPK Thr172 and downstream substrates (P-ACC S79, P-ULK1 S555, P-Raptor S792). Similar results observed in three independent experiments.

**Figure S2.**
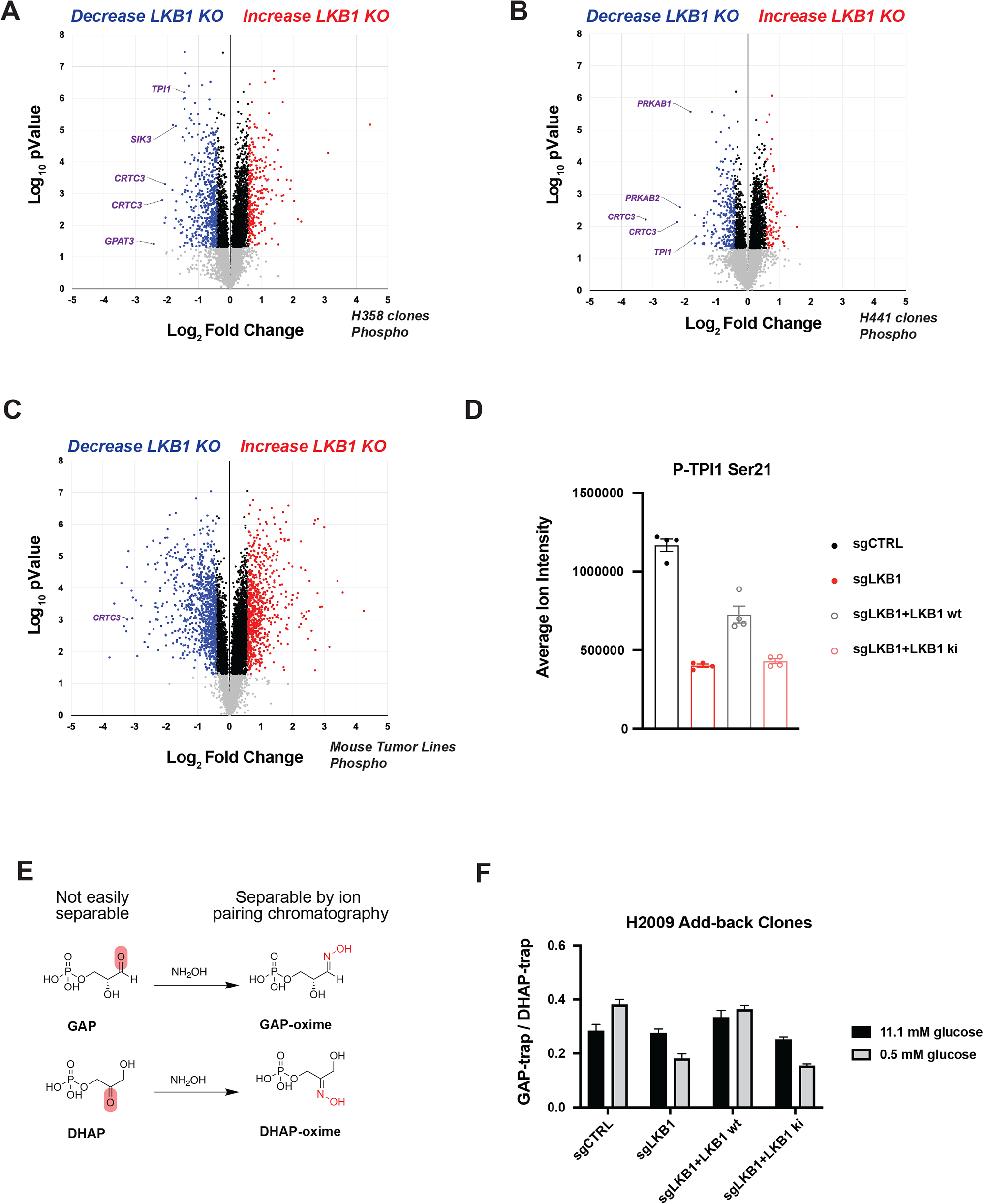
Phosphorylation of human TPI1 is LKB1-dependent and regulates triose phosphate levels. **(A and B)** Volcano plots for comparison of phospho-peptides enriched from lysates of H358 and H441 isogenic clones respectively with and without LKB1 [2 KP clones (H358: sgNT1.4 and sgNT1.6; H441: sgNT1.2 and sgNT1.4) and 2 KPL clones (H358: sgLKB1-2.1 and sgLKB1-3.2. H441: sgLKB1-2.2 and sgLKB1-3.3) with 2 biological replicates for each cell line]. Cells were grown in 0.5 mM glucose for 6 hours before lysis. Phospho-peptides that pass statistical criteria (p-value <0.05) are highlighted in black, red and blue, peptides that do not satisfy this criterion are colored grey. Phospho-peptides highlighted in red satisfy a fold-change threshold (>1.5) upon LKB1 deletion; those highlighted in blue satisfy the fold change threshold (<-1.5) upon LKB1 deletion. Phospho-peptides referenced in the text (SIK3, CRTC3, PRKAB1 and PRKAB2) are labeled in purple text. **(C)** Volcano plot for comparison of quantitative phospho-proteomic data of genetic sensitivity in mLUAD cell-lines, 634T (KP) and Lkb1-t2 (KPL) in biological triplicate for each condition. Analysis conducted on cells treated with 0.5mM glucose for 6 hours in culture. Statistical criteria and color scheme same as for panel A and B. **(D)** Average ion intensity of the H2009 (*KRAS*;*TP53*) isogenic clones (KP: sgNT1.1 and sgNT1.2; KPL: sgLKB1-3.1 and sgLKB1-3.7) and lines with additional transgenic expression of guide RNA resistant LKB1 wildtype (WT) (sgLKB1-3.1 + LKB1 WT and sgLKB1-3.7 + LKB1 WT) or LKB1 kinase inactive (KI) (sgLKB1-3.1 + LKB1 KI and sgLKB1-3.7 LKB1 KI) for the phospho-peptide containing Serine 21 of TPI1 from the experiments from which the volcano plot in Figure 2B was derived. Bar graph depicts each genotype individually and shows restoration of TPI1 phosphorylation in KPL lines expressing transgenic WT LKB1 but not KI LKB1. Ion intensities were normalized to identified non-phosphorylated variant across conditions to control for protein expression; the relevant phospho-peptide was observed 3 times in each biological replicate. **(E)** Schematic showing hydroxylamine chemical labeling and conversion of the triose phosphates; GAP and DHAP to their oxime derivatives. **(F)** *In-situ* chemical trapping metabolomics of hydroxylamine-labeled GAP and DHAP in H2009 clones (KP: sgNT1.1 and sgNT1.2; KPL: sgLKB1-3.1 and sgLKB1-3.7) and additionally lines with transgenic expression of guide RNA resistant LKB1 wildtype (WT) (sgLKB1-3.1 + LKB1 WT and sgLKB1-3.7 + LKB1 WT) or LKB1 kinase inactive (KI) (sgLKB1-3.1 + LKB1 KI and sgLKB1-3.7 LKB1 KI) and treated in culture for 6 hours with 11.1 mM or 0.5 mM respectively. Data presented are representative of three independent biological experiments each containing two technical replicates and reported as the mean ratio (GAP-trap/DHAP-trap) (-/+s.e.m.). Cell number normalized across models 12 hours prior to assay and samples normalized to an exogenous standard, *d* _3_-serine.

**Figure S3.**
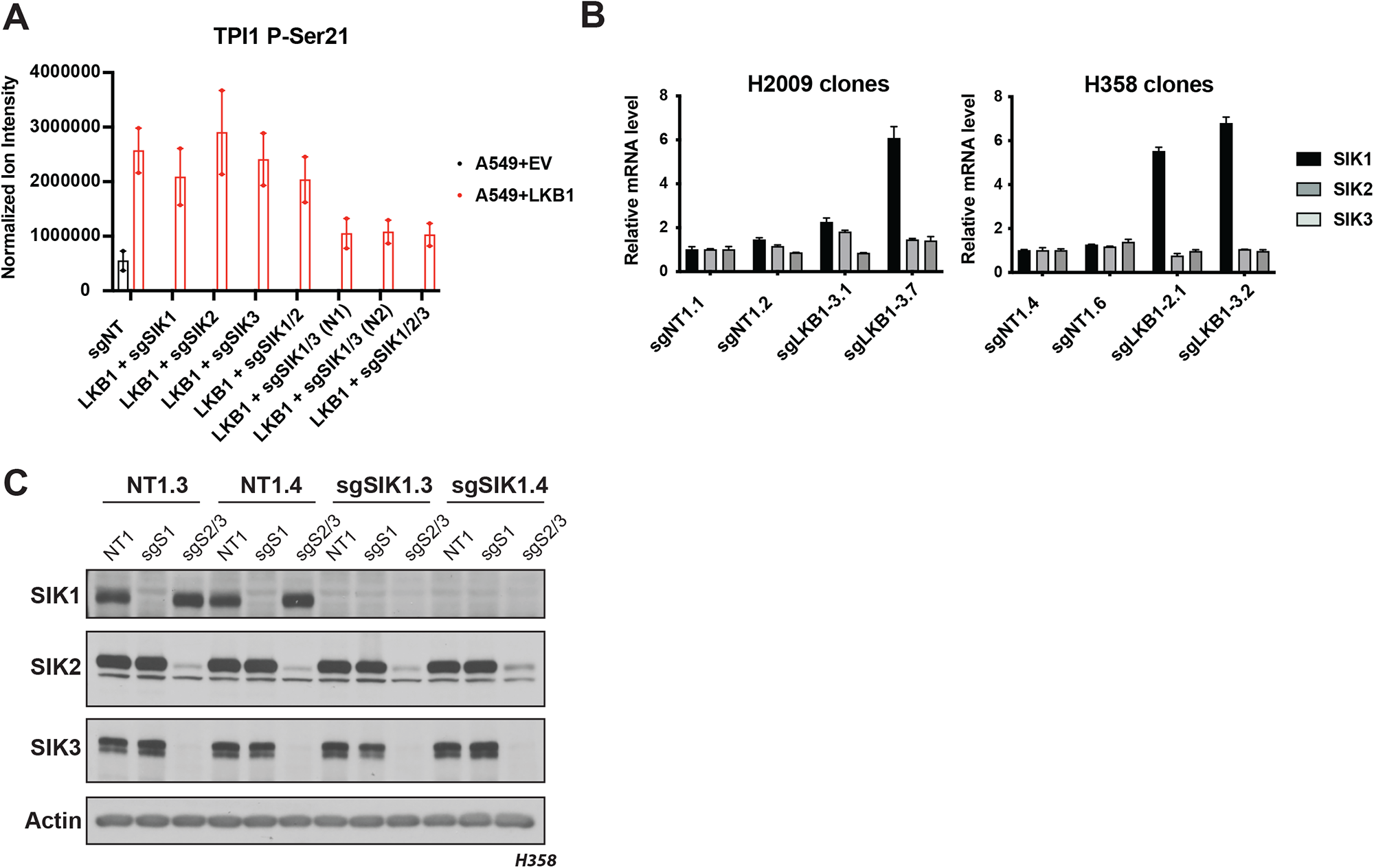
Members of the salt Inducible kinase family phosphorylate TPI1. **(A)** Bar graph of normalized ion abundance for the TPI1-derived Ser-21 phospho-peptide in extracts of A549 cell-lines expressing an empty vector or wild type LKB1 and guide RNAs specifically targeting combinations of members of the Salt Inducible Kinase family. Cell lines were cultured in 11.1 mM glucose. **(B)** Graphs depicting the mean mRNA level (+ s.d.) of the Salt Inducible Kinases in the indicated isogenic clones of H2009 (left) and H358 (right). **(C)** Western blots showing abundance of SIK1, SIK2 and SIK3 in H358 isogenic clones after exposure to the indicated guide RNAs for members of the SIK family. H358 isogenic clones expressing Cas9 and containing non-targeting control (sgNT1.3 and sgNT1.4) or SIK1 specific (sgSIK1.3 and sgSIK1.4) guide RNA and additional non-targeting (NT1), SIK1 (sgSIK1) or dual SIK2 and SIK3 (sgSIK2/3) guide RNAs.

## Methods

No statistical methods were used to predetermine sample size. Data were visualized and statistical analyses performed using Prism 9 software (Graph Pad) or R statistical package. P < 0.05 was considered statistically significant. P values for paired comparisons between two groups with comparable variance were calculated by two-tailed Student’s t-test.

### Reagents

Media and sera were purchased from Life Technologies and R&D Systems respectively. All other reagents were from Sigma-Aldrich unless otherwise noted.

### Cell lines

All cell lines (A549, H358, H441, H2009, 634T and HEK293T) were purchased from ATCC or kindly provided by Kwok-Kin Wong at NYU Langone Medical Center (634T and Lkb1-t2). Cells were maintained in RPMI 1640 medium (Life Technologies: 11879020) supplemented with glucose concentrations as indicated (Life Technologies: A2494001) except for HEK293T cells, which were propagated in DMEM with sodium pyruvate and L-glutamine (Corning). All media supplemented with 10% FBS, 100 units/ml of penicillin and 100 μg/ml streptomycin and grown at 37°C in a 5% CO_2,_ humidified incubator. All cell lines were confirmed to be mycoplasma-free using the MycoAlert mycoplasma detection kit (Lonza: LT07-218).

### CRISPR/Cas9 reagents, plasmids

The control and LKB1-KO lines were generated by infecting the cell lines with lentivirus generated from the LentiCRISPRv2 plasmid (Addgene: 52961). The control and TPI1-KO or SIK-KO lines were generated by infecting the Cas9-expressing lines (LentiCRISPRv2) with lentivirus generated from the LRT2B plasmid (Addgene: 110854). The sgRNA sequences are as follows: sgNT1, CCAATACGGACCGGATTGCT; sgLKB1-2, TGTATAACACATCCACCAGC; sgLKB1-3, TGCACAAGGACATCAAGCCG; sgSAFE, GGTTGGATAAGGCTTAGAAA; sgTPI1-3, GAAGTACACGAGAAGCTCCG; sgTPI1-4, GGAAGCCATCCACATCAGGC; sgSIK1, ATGGTCGTGACAGTACTCCA; sgSIK2, GCACCGGATCACCAAGACGG; sgSIK3, GTGCTTGCAGATCTGCTCCA. The TPI1 alleles were synthesized by Twist Biosciences and cloned into pHAGE-CMV-N-Flag-HA-IRES-Puro-DEST via Gateway cloning.

### Lentivirus production, transduction and single-cell cloning

Lentivirus was generated by transfecting the target plasmid with the packaging plasmids pMD2.G (Addgene: 12259) and psPAX2 (Addgene: 12260) into 293T cells using Lipofectamine 3000 (Invitrogen: L3000015). Media was changed 6 hours after transfection, and then the viral supernatant was collected at 24 and 48h post-transfection. Transduction was conducted in 6 well format on 1 × 10^5^ cells and cells plated in suspension into viral supernatant containing 8 μg/ml Polybrene (Santa Cruz Biotechnology: SC-134220) and incubated overnight (16h). Viral supernatant aspirated and fresh culturing media added to transduced cells for recovery for 24h. Puromycin (Life Technologies: A1113803) was supplemented into media 48h post-transduction for relevant plasmids (LentiCRISPRv2 and pHAGE-CMV-N-Flag-HA-IRES-PURO) at a concentration of 2 μg/ml and selection conducted for 72h. Blasticidin (Invivogen: ANT-BL-1) was supplemented into media 48h post-transduction for relevant plasmid (LRT2B) at a concentration of 10 μg/ml and selection conducted for 5 days. Following selection, single cell cloning was conducted by serial dilution and plating into a 96 well plate, and cells were maintained under relevant selection criteria during the cloning process. Clones that grow out from single cells were expanded and validation of knockout conducted by western-blot or qPCR as indicated.

### LUAD clinical data set analysis

Human LUAD (hLUAD) datasets (TCGA and MSKCC) were downloaded from cBioPortal and KRAS, TP53, and LKB1 mutational and copy number status were assessed. Samples were divided into KRAS-mutant and KRAS-wild-type cohorts for further analysis. Using the R statistical software package, a Fisher’s exact test was performed on each cohort to determine the odds of TP53 and LKB1 mutations co-occurring.

### Mice and xenografts

Animal procedures were performed with the approval of the Weill Cornell Medicine IACUC. Tumor volume was not allowed to exceed 1000 mm^3^ in any experiment. Prior to implantation, cells were re-suspended in PBS and mixed 1:1 with Matrigel (Corning, 356231). Cells were then injected subcutaneously into single flanks of 6-week-old athymic mice (Envigo). Caliper measurements were performed weekly to monitor tumor growth. For the H358 LKB1-KO clones, 1 × 10^6^ cells were injected per flank; for the murine lung tumor lines (634T and Lkb1-t2) 1 × 10^4^ cells were injected per flank.

### Western blotting

Protein lysates were prepared in CST lysis buffer (Cell Signaling Technology: 9803) supplemented with cOmplete mini EDTA free protease inhibitor (Roche: 04693159001) and quantified using the BCA protein assay (Thermo Scientific, 23225). Lysates prepared at a concentration of 1 mg/ml and supplemented with 4x Laemmli Sample Buffer (Bio-Rad: 1610747) supplemented with fresh 2-mercaptoethanol (Sigma: M3148). Proteins were separated on self-cast Tris-glycine polyacrylamide gels, transferred to Polyscreen PVDF membranes (Perkin Elmer: NEF1002), and probed with Cell Signaling Technology antibodies used at 1:1000 in 5% BSA (Sigma: A4503) in TBS-T: P-ACC Ser79 (#3661), ACC (#3662), P-Raptor Ser792 (#2083), Raptor (#2280), P-AMPKα Thr172 (#2535), AMPK α1/2 (#2532), LKB1 (#3047), P-ULK1 Ser555 (#5869), ULK1 (#8054) and SIK2 (6919). Antibodies from Abcam used at concentrations indicated in 5% BSA in TBS-T against β-actin (ab6276, 1:20,000), TPI1 (ab96696, 1:3,000) and P-SIK1 Thr182 + P-SIK2 Thr175 + P-SIK3 Thr163 (ab199474, 1:1000). Antibody from Novus Biologicals was used at 1:20,000 in 5% BSA in TBS-T against SIK3 (NBP2-47278). Antibodies from Sigma-Aldrich against Flag epitope tag (F7425, 1:5,000) and (F3165, 1:1000) were used at indicated concentrations in 5% BSA in TBS-T. Secondary antibodies from Millipore against Rabbit (AP132PMI) and Mouse (AP124PMI) primary antibodies were resuspended per manufacturer’s instructions and used at 1:10,000 in 5% non-fat dried milk in TBS-T. Western blots were then developed in the dark room on an autoradiograph following incubation with home-made ECL.

### Sn-glycerol-3-phosphate steady state analysis metabolite extraction

Cells were cultured in medium reconstituted from glucose-free RPMI 1640 medium (Life Technologies: 11879020) supplemented with 11.1 or 0.5 mM glucose and 10% dialyzed FBS. The day prior to treatment and collection cells were lifted and counted and 2 × 10^6^ cells were plated in a 10cm culture dish. Cells were given a medium change 1 h before the addition of growth medium. Cells were rinsed twice with PBS before the addition of tracing medium. The time of addition of tracer medium was designated time 0. Metabolites were extracted at 30 minutes post addition as indicated in text.

### Aqueous metabolite extraction and liquid chromatography–mass spectrometry (LC–MS) analysis

Cells were washed twice with PBS, and twice with LC-MS grade H_2_O. Five hundred μl of 80% methanol at −80 °C was added to quench metabolic reactions and the cells were collected by scraping. The lysate was then transferred to a fresh 2.0 ml Eppendorf tube pre-chilled on dry-ice and an additional 500 μl of 80% methanol was added to the original plate and scraped again. The second lysate was added to the first and incubated on dry ice for 20 minutes with intermittent vortexing then centrifuged at 16,000g for 10 min to allow cellular debris to be pelleted. The aqueous volume was then transferred to a clean, fresh pre-chilled 2.0 ml Eppendorf tube and dried under vacuum in a speedvac and stored at −80°C. Dried sample pellets were resuspended in HPLC-grade water (20 μl) and centrifuged at 20,000 g for 5 min to remove insoluble material. Following centrifugation, 16 μl of supernatant was transferred to virgin polypropylene auto sampler vials, capped and placed on dry ice. Supernatants (5 μl) were injected and analyzed using a hybrid 6500 QTRAP triple quadrupole mass spectrometer (AB/SCIEX) coupled to a Prominence UFLC HPLC system (Shimadzu) via selected reaction monitoring (MRM). ESI voltage was +4900V in positive ion mode with a dwell time of 3 ms per SRM transition. Approximately 10–14 data points were acquired per detected metabolite. Samples were delivered to the mass spectrometer via hydrophilic interaction chromatography (HILIC) using a 4.6 mm i.d. x 10 cm Amide XBridge column (Waters) at 400 μl/min. Gradients were run starting from 85% buffer B (HPLC grade acetonitrile) to 42% B from 0 to 5 min; 42% B to 0% B from 5 to 16 min; 0% B was held from 16 to 24 min; 0% B to 85% B from 24 to 25 min; 85% B was held for 7 min to re-equilibrate the column. Buffer A was comprised of 20 mM ammonium hydroxide/20 mM ammonium acetate (pH = 9.0) in 95:5 water:acetonitrile. Peak areas from the total ion current for each metabolite SRM transition were integrated using MultiQuant v3.0 software (AB/SCIEX). Tubes containing cellular debris was retained to determine protein concentration for data normalization. Briefly pellet was resuspended by addition of 600 μl of sodium hydroxide and boiled at 90 °C for 30 minutes with intermittent vortexing. Resolubilized pellets were allowed to come to room temperature, and protein quantified using the DC protein assay (Bio-Rad: 5000111). Derived metabolite data was normalized to protein concentration and median ion intensity per injection across the dataset.

### *In situ* hydroxylamine trapping in live cells

Two 15 cm dishes per condition were plated with 9 × 10^6^ cells 24 hr prior to treatment. Plated cells were washed twice with PBS then grown in RPMI 1640 media containing 11.1 mM or 0.5 mM glucose as indicated for 6 hr. Cells were then washed twice with PBS and 3 ml of PBS containing protease inhibitors was added to the plate and cells were scraped. Cell homogenate was transferred to a 15ml conical tube and centrifuged at 1,400 x g for 3 minutes to pellet cells. Cellular pellets were resuspended in 300 μl ice cold 80% Methanol and transferred to a 1.5 ml Eppendorf tube. Chemical labeling of live cells was achieved by adding 10 μl of hydroxylamine solution (Sigma: 467804, ~15M solution) and incubated for 10 minutes with gently vortexing intermittently. Following a 10 minute incubation, the suspended cells were lysed with a probe sonicator set to 30% amperage pulse (1:1 pulse:pause 16 seconds total). Lysed cellular homogenates were then centrifuged at 20,000 x g for 10 minutes at 4 °C. Clarified supernatant was transferred to a fresh 1.5 ml Eppendorf tube and dried under Nitrogen gas flow until all solvent was evaporated. Dried pellets were then stored at −80 °C until ready for analysis. Dried metabolites were resuspended in 100 μL of an 80:20 mixture of MeOH/H_2_O and an internal deuterated standard, 10 nmol *d*_3_*-*serine, was added to the dried metabolome solution for quantification and sample normalization.

### Targeted LC-MS/MS for hydroxylamine trapping

Resuspended metabolites were separated by hydrophilic interaction chromatography with a Gemini reverse-phase C18 column (50 mm × 4.6 mm with 5 μm diameter particles) from Phenomenex together with precolumn (C18, 3.5 mm, 2 mm × 20 mm). Mobile phase A was composed of 100% H_2_O (10 mM tributylamine aqueous solution, adjusted to pH 4.95 with 15 mM acetic acid), and mobile phase B was composed of 100% Methanol. Using a multi-step gradient with buffer A and B: 0-5 min, 95% A; 5-15 min, 95-90% A; 15-22 min, 90-85% A; 22-26 min, 10% A, and maintained for 4 min; 30-33 min, 95% A, and maintained for 7 min. The flow rate was 0.2 ml/min for 0-15 min and 30-40 min, and 0.3 ml/min for 15-30 min. Targeted MS/MS analysis was performed on an Agilent triple quadrupole LC-MS/MS instrument (Agilent Technologies 6460 QQQ). The capillary voltage was set to 4.0 kV. The drying gas temperature was 350°C, the drying gas flow rate was 10 L/min, and the nebulizer pressure was 45 psi. Relative metabolite abundance was quantified by integrated peak area for the given MRM-transition. Data presented are representative of three independent biological experiments each containing three technical replicates for a given condition.

### Proteomics and phospho-proteomic sample preparation

Protein lysates were prepared in CST lysis buffer (Cell Signaling Technology: 9803) supplemented with cOmplete mini EDTA free protease inhibitor (Roche: 04693159001) and quantified using the BCA protein assay (Thermo Scientific, 23225). Following quantification, 100 μg of each protein lysate was moved into a clean 1.5 mL tube. Following distribution of protein, each tube was brought to a final volume of 300 μL by addition of PBS, followed by precipitation with trichloroacetic acid (TCA) (Sigma) to a final concentration of 25%, vigorously vortexed and incubated on ice overnight. TCA precipitates were centrifuged at 21,130 *x g* for 30 minutes at 4°C, washed twice in 500 µL of ice-cold acetone, and centrifuged at 21,130 *x g* for 10 minutes after each wash. Following precipitation and washes, pellets were allowed to completely dry at room temperature. Dry pellets were re-suspended in 100 μL of 100 mM TEAB, 0.5% SDS and reduced with 9.5 mM tris-carboxyethyl phosphine (TCEP) for 60 minutes at 55°C. Following reduction of disulfide bonds with TCEP, the denatured protein mix was centrifuged at 21,130 *x g* for 5 minutes then alkylated with 4.5 mM iodoacetamide (IA) for 30 minutes in the dark at room temperature. After reduction and alkylation of disulfide bonds, the denatured protein mixture was precipitated out of solution by addition of 600 µL of ice-cold acetone and placed in the −20°C freezer overnight. The following day precipitated proteins were centrifuged at 8,000 *x g* for 10 minutes to pellet precipitated protein. Following centrifugation supernatant was decanted off and pellets were allowed to air-dry at room temperature. Once dry, protein pellets were reconstituted in 100 µL 100 mM TEAB and CaCl_2_ was supplemented to a final concentration of 1 mM, 2 µg of sequencing grade Trypsin (Promega) was added, and reactions were placed in the dark on a thermal mixer (Eppendorf) set to 37°C and shaking at 850 r.p.m. for 16 hours.

### Thermal Proteomic Profiling

Cells were lifted using TrypLE Express (Thermo Fisher Scientific - GIBCO) and neutralized following 5-minute incubation using complete media (RPMI + 10% FBS penicillin/streptomycin) and centrifuged at 1100 r.p.m. for 4 minutes. The cell pellet was reconstituted in 10 mL PBS containing protease and phosphatase inhibitors (Roche) and centrifuged again at 1100 RPM for 4 minutes. Following centrifugation, the cell pellet was resuspended in 1 mL PBS with inhibitors and distributed into thin-wall PCR tubes at 100 μL of cell suspension in each tube. Thermal denaturation was performed as previously described, and the resulting cellular suspension was transferred to clean 1.5 mL microcentrifuge tubes and PCR tubes were additionally rinsed with 30 μL of PBS with inhibitors to ensure complete transfer of cellular suspension. Cellular suspension was next snap frozen in liquid nitrogen for 1 minute followed by thawing and re-equilibration back to room temperature. This freeze-thaw cycle was repeated 2 additional times and the soluble fraction of each lysate was generated by centrifugation at 21,130 *x g* for 30 minutes at 4°C. Supernatants were transferred to clean 1.5 mL microcentrifuge tubes, and protein was quantified in the supernatant for temperatures 37°C and 41°C by micro-BCA assay (Thermo Fisher Scientific - Pierce). Following quantification, the average of the two lowest temperatures was taken and the volume equivalent to 30 μg of protein in the lowest temperature was moved from each temperature fraction into a clean 1.5 mL tube. Following distribution of protein, each tube was brought to a final volume of 300 μL by addition of PBS with inhibitors, followed by precipitation with trichloroacetic acid (TCA) (Sigma) to a final concentration of 25%, vigorously vortexed and incubated on ice overnight. TCA precipitates were centrifuged at 21,130 *x g* for 30 minutes at 4°C, washed twice in 500 µL of ice-cold acetone, and centrifuged at 21,130 *x g* for 10 minutes after each wash. Following precipitation and washes, pellets were allowed to completely dry at room temperature. Dry pellets were re-suspended in 100 μL of 100 mM TEAB, 0.5% SDS and reduced with 9.5 mM tris-carboxyethyl phosphine (TCEP) for 60 minutes at 55°C. Following reduction of disulfide bonds with TCEP, the denatured protein mix was centrifuged at 21,130 *x g* for 5 minutes then alkylated with 4.5 mM iodoacetamide (IA) for 30 minutes in the dark at room temperature. After reduction and alkylation of disulfide bonds, the denatured protein mixture was precipitated out of solution by addition of 600 µL of ice-cold acetone and placed in the −20°C freezer overnight. The following day precipitated proteins were centrifuged at 8,000 *x g* for 10 minutes to pellet precipitated protein. Following centrifugation supernatant was decanted off and pellets were allowed to air-dry at room temperature. Once dry, protein pellets were reconstituted in 100 µL 100 mM TEAB and CaCl_2_ was supplemented to a final concentration of 1 mM, 1 µg of sequencing grade Trypsin (Promega) was added, and reactions were placed in the dark on a thermal mixer (Eppendorf) set to 37°C and shaking at 850 r.p.m. for 16 hours. The next day, digested samples were centrifuged at 21,130 *x g* for 10 minutes and proceeded to TMT labeling of digested samples.

### TMT Labeling, Fractionation, and Phosphopeptide Enrichment

TMT labeling was performed generally as per manufacturer’s protocol. Briefly, each TMT tag was re-suspended in 164 μL anhydrous acetonitrile with intermittent vortexing for 10 minutes. Following resuspension, 41 μL was added to corresponding temperatures (TMT-126 = 37°C; four separate aliquots of each temperature for subsequent desalting and fractionation) and labeling reaction was allowed to proceed for 1 hour at room temperature. Reactions were quenched by addition of 8 μL of 5% hydroxylamine in 100 mM TEAB and incubated for 15 minutes. Labeled temperature fractions were pooled, desalted on 1cc/50 mg C18 SepPAK columns (Waters # WAT054955) on a vacuum manifold and desalted peptides were dried down in a speedvac. Dried peptides were reconstituted in 300 µL of 0.1% TFA in H_2_O, high-pH reverse phase spin-columns (Thermo fisher scientific - Pierce) were equilibrated, and samples fractionated per manufacturer’s instructions into 8 fractions, 2 washes and a flow-through fraction (11 total). Separate samples from the same fractions were then combined and dried. Peptide fractions were reconstituted in 200 µL of 5% acetonitrile, 0.1% TFA in water, and 10 µL was removed for bulk HTP analysis. The remaining fractionated labeled peptides dried and re-dissolved in 40% acetonitrile, 6% TFA in water before phosphopeptide enrichment with Titansphere 5 µm TiO_2_ beads (GL Sciences). Titansphere TiO2 beads (GL Sciences) were reconstituted in buffer containing 80% acetonitrile, 6% TFA, and 2,5-dihydroxybenzoic acid (20 mg/mL) and rotated for 15 min at 25°C. Equal amount of beads slurry (~5:1 beads-to-peptide ratio based on concentration of peptides in 37°C aliquot) was added to each temperature aliquot of reconstituted peptides and rotated for 20 mins 25°C. Beads were then washed twice with higher percentage of acetonitrile (10% and 40%) in 6% TFA and supernatant was removed by centrifugation at 500 *x g* for 2 min. Washed beads were then added to self-packed stage tip with C8 SPE (Sigma Aldrich) and washed once more with 60% acetonitrile in 6% TFA. Phosphopeptides were first eluted with 5% NH_4_OH, then 10% NH_4_OH, 25% acetonitrile, and dried with speedvac. Dried phosphopeptides were reconstituted in 5% acetonitrile, 1% TFA, desalted with self-packed stage tip with C18 SPE (Sigma Aldrich), and dried with speedvac once more. The final processed phosphopeptides were reconstituted in 5% acetonitrile, 0.1% TFA in water for LC-MS^3^ analysis.

### LC-MS^3^ Analysis and Data Acquisition

High-pH reverse-phase fractions were run on a 4-hour instrument method with an effective linear gradient of 180 minutes from 5% to 25% mobile phase B with the following mobile phases: A: 0.1% formic acid in H_2_O, B: 80% acetonitrile/0.1% formic acid in water on a 50 cm Acclaim PepMap RSLC C18 column (Thermo Fisher Scientific #164942) operated by a Dionex ultimate 3000 RSLC nano pump with column heating at 50°C connected to an Orbitrap Fusion Lumos. Briefly, the instrument method was a data-dependent analysis and cycle time set to 3 seconds, total. Each cycle consisted of one full-scan mass spectrum (400-1500 m/z) at a resolution of 120,000, RF Lens: 60%, maximum injection time of 100 ms followed by data-dependent MS/MS spectra with precursor selection determined by the following parameters: AGC Target of 4.0e^5^, maximum injection time of 100 ms, monoisotopic peak determination: peptide, charge state inclusion: 2-7, dynamic exclusion 10 sec with an intensity threshold filter: 5.0e^3^. Data-dependent MS/MS spectra were generated by isolating in the quadrupole with an isolation window of 0.4 *m*/z with CID activation and corresponding collision energy of 35%, CID activation time of 10 ms, activation Q of 0.25, detector type Ion Trap in Turbo mode, AGC target of 1.0e4 and maximum injection time of 120 ms. Data-dependent multi-notch MS^3^ was done in synchronous precursor selection mode (SPS, multi-notch MS^3^) with the following settings: Precursor selection Range; Mass Range 400-1200, Precursor Ion Exclusion Properties *m*/z Low: 18 High: 5, Isobaric Tag Loss Exclusion Properties: TMT. Number of SPS precursors was set to 10 and data-dependent MS^3^ was detected in the Orbitrap (60,000 resolution, scan range 120-500) with an isolation window of 2 *m*/z HCD activation type with collision energy of 55%, AGC target of 1.2e5 and a maximum injection time of 150 ms. Raw files were parsed into MS1, MS2 and MS3 spectra using RawConverter.

### Proteomic, phospho-proteomic and Thermal Profiling Data Analysis

Data generated were searched using the ProLuCID algorithm in the Integrated Proteomics Pipeline (IP2) software platform. Human and Mouse proteome data were searched using concatenated target/decoy UniProt databases. Basic searches were performed with the following search parameters: HCD fragmentation method; monoisotopic precursor ions; high resolution mode (3 isotopic peaks); precursor mass range 600-6,000 and initial fragment tolerance at 600 p.p.m.; enzyme cleavage specificity at C-terminal lysine and arginine residues with 3 missed cleavage sites permitted; static modification of +57.02146 on cysteine (carboxyamidomethylation), +229.1629 on N-terminal and lysine for TMT-10-plex tag; 4 total differential modification sites per peptide, including oxidized methionine (+15.9949), and phosphorylation (+79.9663) on serine, threonine, and tyrosine (only for phospho-enriched samples); primary scoring type by XCorr and secondary by Zscore; minimum peptide length of six residues with a candidate peptide threshold of 500. A minimum of one peptide per protein and half-tryptic peptide specificity were required. Starting statistics were performed with a Δmass cutoff = 10 p.p.m. with modstat, and trypstat settings. False-discovery rates of protein (pfp) were set to 1% (for unenriched datasets) or peptide (sfp) set to 1% (for phospho-proteomics datasets). TMT quantification was performed using the isobaric labeling 10-plex labeling algorithm, with a mass tolerance of 5.0 p.p.m. or less. Reporter ions 126.127726, 127.124761, 127.131081, 128.128116, 128.134436, 129.131417, 129.13779, 130.134825, 130.141145, and 131.13838 were used for relative quantification.

